# ALDH1-positive intratumoral stromal cells indicate epithelial differentiation and good prognosis in prostate cancer

**DOI:** 10.1101/276378

**Authors:** Paulina Nastały, Martyna Filipska, Colm Morrissey, Elke Eltze, Axel Semjonow, Burkhard Brandt, Klaus Pantel, Natalia Bednarz-Knoll

## Abstract

Aldehyde dehydrogenase 1 (ALDH1) characterizes tumor-initiating cells in solid tumors, however little is known about its expression in intratumoral stromal cells. Herein, we aimed to dissect its potential dual relevance in prostate cancer (PCa).

ALDH1 expression was evaluated immunohistochemically in tumor and stromal cells in primary PCa and metastasis. It was correlated with clinico-pathological parameters, outcome of patients, and selected protein expression (CK5/6, CK14, CK8/18, CK19, EpCAM, Ki-67, E-cadherin, N-cadherin, and vimentin).

ALDH1 protein was detected in tumor and stromal cells in 16% and 67% of 348 primary PCa, respectively. Tumor cell ALDH1 expression was associated with advanced tumor (T) status (p=0.009), higher Gleason score (p=0.016), shorter time to biochemical recurrence (BR) (p=0.010) and CK14 expression (p=0.023). Stromal cell ALDH1 expression correlated with lower T status (p=0.008), N0 status (p=0.017), lower Gleason score (p=0.016) and longer time to BR (p=0.017). In the subgroup of d’Amico high-risk patients it occurred even to be an independent predictor of good prognosis (multivariate analysis, p=0.050). ALDH1 was found in stroma of tumors characterized by CK8/18 (p=0.033) or EpCAM expression (p<0.001) and rarely by epithelial-mesenchymal transition defined as CK8/18(-)vimentin(+) phenotype (p=0.003). ALDH1 was detected in tumor cells and stroma of 33% and 41% of hormone naive lymph node metastases (n=63), 52% and 24% of castration resistant bone metastases, as well as 89% and 28% of castration resistant visceral metastases (n=21), respectively.

We have determined that contrary to tumor cell ALDH1, the presence of stromal ALDH1 is associated with a more differentiated tumor epithelial phenotype in primary PCa, improved clinical outcome, and is less frequent in PCa metastases.

**Abbreviations:**

ALDH1: aldehyde dehydrogenase 1
BR: biochemical recurrence
CK: cytokeratin
CRPC: castration resistant prostate cancer
CSC: cancer stem cell
FFPE: formalin fixed paraffin embedded tissue
IHC: immunohistochemistry
LNM: lymph node metastasis
OS: overall survival
PSA: prostate specific antigen
PT: primary tumor
REMARK: REporting recommendations for tumour MARKer prognostic studies
TMA: tissue microarray

## Introduction

It has been suggested that aldehyde dehydogenase 1 (ALDH1) in breast cancer might play an opposing role depending on the type of cells, in which it is expressed (1, 2). The presence of ALDH1 in tumor cells of breast cancer and other solid tumors was reported to define a population of ‘so called’ cancer stem cells (CSC) or tumor progenitor cells (3). Such cells are believed to initiate and drive tumor progression and represent a subpopulation of cells particularly resistant to applied therapies (4). Accordingly, tumor cell ALDH1 staining is associated with a more aggressive disease course in solid tumors (5). On the contrary, the presence of ALDH1 in tumor-associated stroma correlates to patients’ better outcome in breast carcinomas (1, 2), although the mechanism of this phenomenon remains largely unknown. ALDH1 is also known to be active in the late steps of retinoic acid synthesis (6), a well-known inducer of epithelial cell differentiation and an inhibitor of proliferation and migration (7, 8, 9). Its involvement in retinoic acid metabolism was demonstrated to mediate differentiation of normal human mammary epithelium (10). Hypothetically, the role of ALDH1 in this process might be cell-specific and stromal cells expressing ALDH1 might modulate tumor development via production of retinoic acid (2). This phenomenon highlights the importance of the interactions between tumor cells and their microenvironment.

Most frequently investigated in breast cancer (11), ALDH1 has been detected also in tumor cells of prostate cancer (PCa) (12) where it was shown to be associated with worse clinical outcome (13, 14). ALDH1-positive cells were shown to be potential tumor-initiating and metastasis-initiating cells in human PCa (15). Additionally, ALDH1-expressing cells might contribute to tumor resistance to radiotherapy (16). To the best of our knowledge, stromal ALDH1 staining has never been reported in PCa. Therefore, in the current study, ALDH1 was investigated as a protein of potential dual clinical and biological relevance in PCa. ALDH1 expression was assessed in PCa cells as well as intratumoral stromal cells in primary PCa and unmatched PCa metastases. The results were compared to clinico-pathological data as well as patients’ outcome. Moreover, in order to test impact of the presence of cell-specific ALDH1 on tumor cell differentiation and aggressiveness, the results were analysed in relation to different cytokeratins (CK5/6, CK14, CK8/18, CK19), a proliferation marker (Ki-67), EpCAM, and epithelial mesenchymal transition-related markers (E-cadherin, N-cadherin, vimentin) in primary PCa.

## Material and methods

### Patients

Three-hundred-ninety-eight patients with sporadic PCa were included in this study based on their signed informed consent form. The patients underwent radical prostatectomy at the Department of Urology in Prostate Centre University Clinic Münster (Germany) during 1998-2003. The variable clinico-pathological (including TNM status according to AJCC Cancer Staging 2010) and molecular parameters were documented as described (ref: 17, 18; **Suppl. Tab. 1**). Biochemical progression during follow-up was defined as a serum prostate specific antigen (PSA) level increasing above 0.1 ng/mL in two consecutive determinations. The timepoint of biochemical progression was considered to be the median time between the last PSA <= 0.1 ng/mL and the first PSA > 0.1 ng/mL. Last follow-up was completed in October 2014. The median follow-up for this cohort of patients was 54 months (range 0.2-176 months).

In addition, 98 hormone naive PCa patients from the Prostate Centre University Clinic Muenster (Germany) were selected for examination of their metastases to lymph nodes.

Moreover, visceral and bone metastases were obtained from 21 PCa patients who died of metastatic castration resistant PCa (CRPC) and who signed written informed consent for a rapid autopsy performed within 6 hours of death, under the aegis of the Prostate Cancer Donor Program at the University of Washington and approved by the Institutional Review Board of the University of Washington. The study was conducted according to REMARK study recommendations (19).

### TMA

Two tissue microarrays (TMAs) with primary PCa samples were prepared as described (17). Briefly, each TMA comprised of 0.6 mm-diameter tissue cores obtained from formalin-fixed paraffin embedded PCa specimens. Fragments of normal prostate and kidney tissues were introduced to TMAs as internal controls. All patients were represented by duplex tumor samples (in case of multifocal disease originated from two different tumor foci).

One-hundred-ninety-six samples of LNM from 98 hormone naive PCa patients were used to construct the LNM tissue microarray (TMA). Within the LNM TMA each patient was represented by duplicate cores.

Eighty-four CRPC metastases from 21 rapid autopsy patients (up to 4 sites per patient) were fixed in buffered formalin (bone metastases were decalcified in 10% formic acid) and embedded in paraffin and were used to construct the CRPC TMA using duplicate 1 mm diameter cores from these tissues.

All TMA sections were cut 4-µm-thick and placed on charged polylysine-coated slides (Superfrost Plus, BDH, Germany) for further examination.

### Immunohistochemistry and its evaluation

ALDH1 staining and its evaluation was performed as described (2). Briefly, mouse monoclonal anti-ALDH1 antibody (44/ALDH1, BD Biosciences, US) diluted 1:500 was incubated overnight at 4°C and envisioned by DAKO ChemMate Detection Kit Peroxidase/DAB, Rabbit/Mouse (Dako, Denmark) to detect ALDH1 protein in PCa. For ALDH1 staining in tumor cells, intensity of the staining (negative, weak, moderate or strong) was multiplied by percentage of the stained tumor cells to result in index score of 0 to 300. The mean value of all index scores was used as a cut-off to determine negative (index score lower than mean) or positive expression (index score equal or greater than mean). ALDH1 expression in stromal cells was determined as no expression, moderate or strong expression in less than 10%, in 10-50% and more than 50% of stromal cells. The results were classified as negative or positive stromal staining according to the cut-off equal 10% to positive cells. Immunohistochemistry for CK5/6, CK14, CK8-18, CK19, E-cadherin, N-cadherin, vimentin, EpCAM and Ki-67 were performed as described (18).

### Statistics

Statistical analysis was performed using SPSS software licensed for Medical University of Gdańsk. Chi-square, and Fisher’s exact tests, as well as Pearson two-tailed correlation test and independent samples t-test were used in order to compare the results to molecular factors and clinico-pathological parameters. Associations between protein expression profiles and time to biochemical recurrence and metastasis-free survival were evaluated using Log Rank (Mantel Cox) test and Kaplan-Meier plot. To estimate hazard risk, Cox-Hazard-Potential regression analysis (CI 95%) was done. All results were considered statistically significant if p<0.05 and highly statistically significant if p<0.001.

## Results

### ALDH1 expression in tumor and stromal cells of primary prostate cancers

Five-hundred-fifty-one tumor samples from 348 PCa patients were informative for ALDH1 staining in tumor and stromal cells.

Index score (i.e. multiplication of the staining intensity by percentage of the stained tumor cells) was used to assess ALDH1 expression in tumor cells. The mean tumor ALDH1 index score was 17.11. Fifty-five (15.8%) patients were positive for ALDH1 staining in tumor cells based on ALDH1 index score categorized according to this mean as cut-off. The detected ALDH1 staining was localized in the cytoplasm of the cancer cells (**Fig. 1**). The percentage of tumor cells positive for ALDH1 ranged from 1 to 99% per tumor sample with the mean of 5.9%.

**Figure 1.**
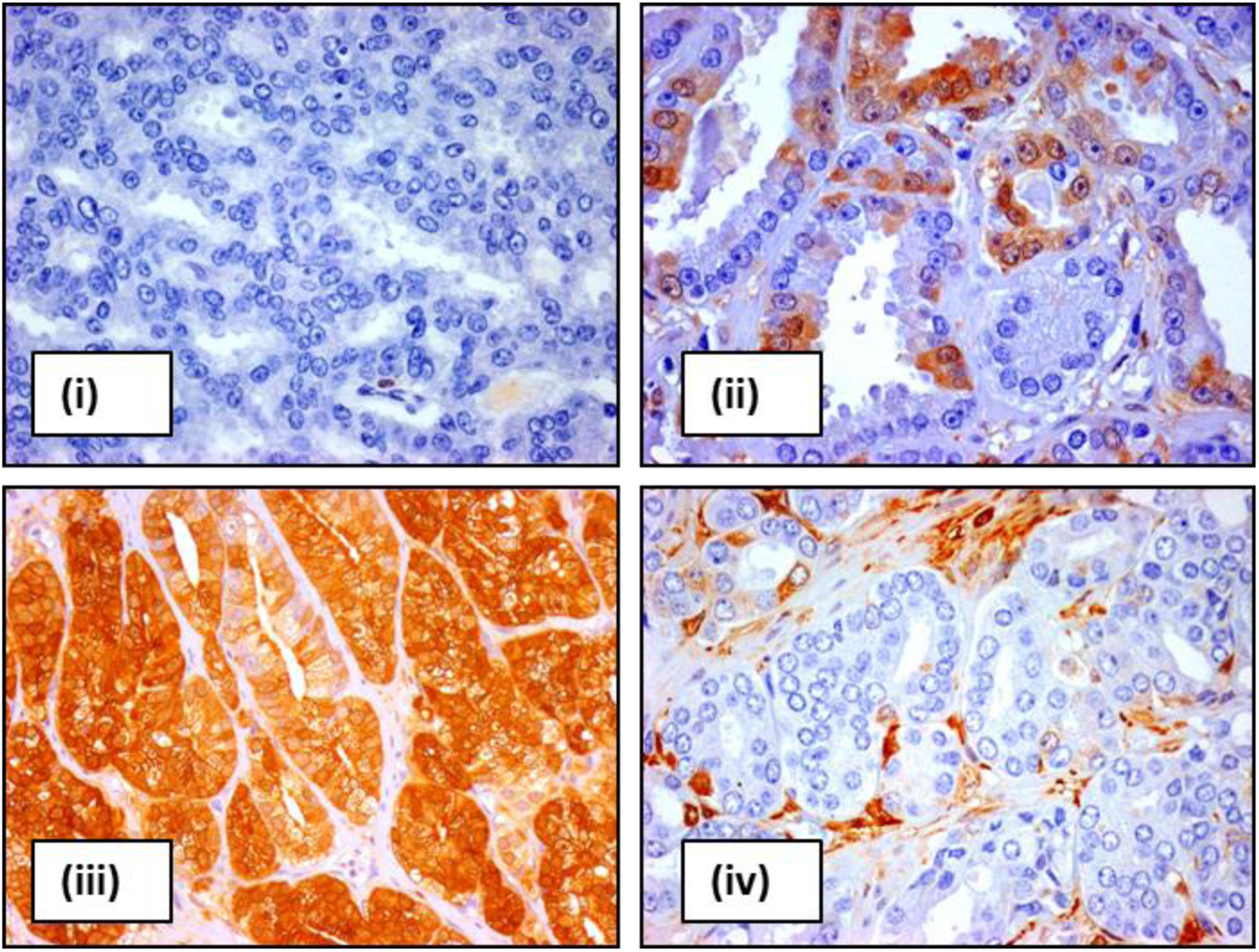
ALDH1 expression in tumor and stromal cells of prostate cancer patients. Representative pictures of ALDH1 staining in prostate cancer samples: no (i, iv); low (ii) and high percentage of ALDH1-positive tumor cells (iii) as well as no (i-iii) and high percentage of ALDH1-positive stromal cells (iv).

Positive ALDH1 stromal expression was found in 234 (67.2%) patients. If present, ALDH1 was detected as moderate or strong cytoplasmic staining in spindle- and/or polygonal-like shaped stromal cells located between or around tumor cells (**Fig. 1**).

Positive ALDH1 staining in exclusively stromal cells was observed in 62.0%, whereas only in tumor cells in 3.2% of patients. ALDH1 positivity in both tumor and stromal cells was found in 7.2% of cases. Lack of both ALDH1 stromal and tumoral expression was seen in 27.6% of PCa.

### Associations of ALDH1 expression in primary prostate cancers to clinico-pathological parameters and patients’ outcome

ALDH1 positive tumor cell expression was more frequent in primary PCa cases with more advanced T status (Chi^2^=6.781, p=0.009), and biochemical recurrence (BR) (Chi^2^=7.808, p=0.005) (**Tab. 1**). ALDH1 tumor cell positivity also displayed a borderline correlation to metastatic relapse (Fisher’s exact test, p=0.058) and a statistically significant association with PCa-related death (Fisher’s exact test, p=0.025) although the number of positive cases was very low in this comparison (**Tab. 1**). Patients positive for ALDH1 in tumor cells had shorter time to BR (Kaplan-Meier log rank analysis, p=0.010) and showed a trend towards shorter time to metastasis (Kaplan-Meier log rank analysis, p=0.051) (**Fig. 2**).

**Figure 2.**
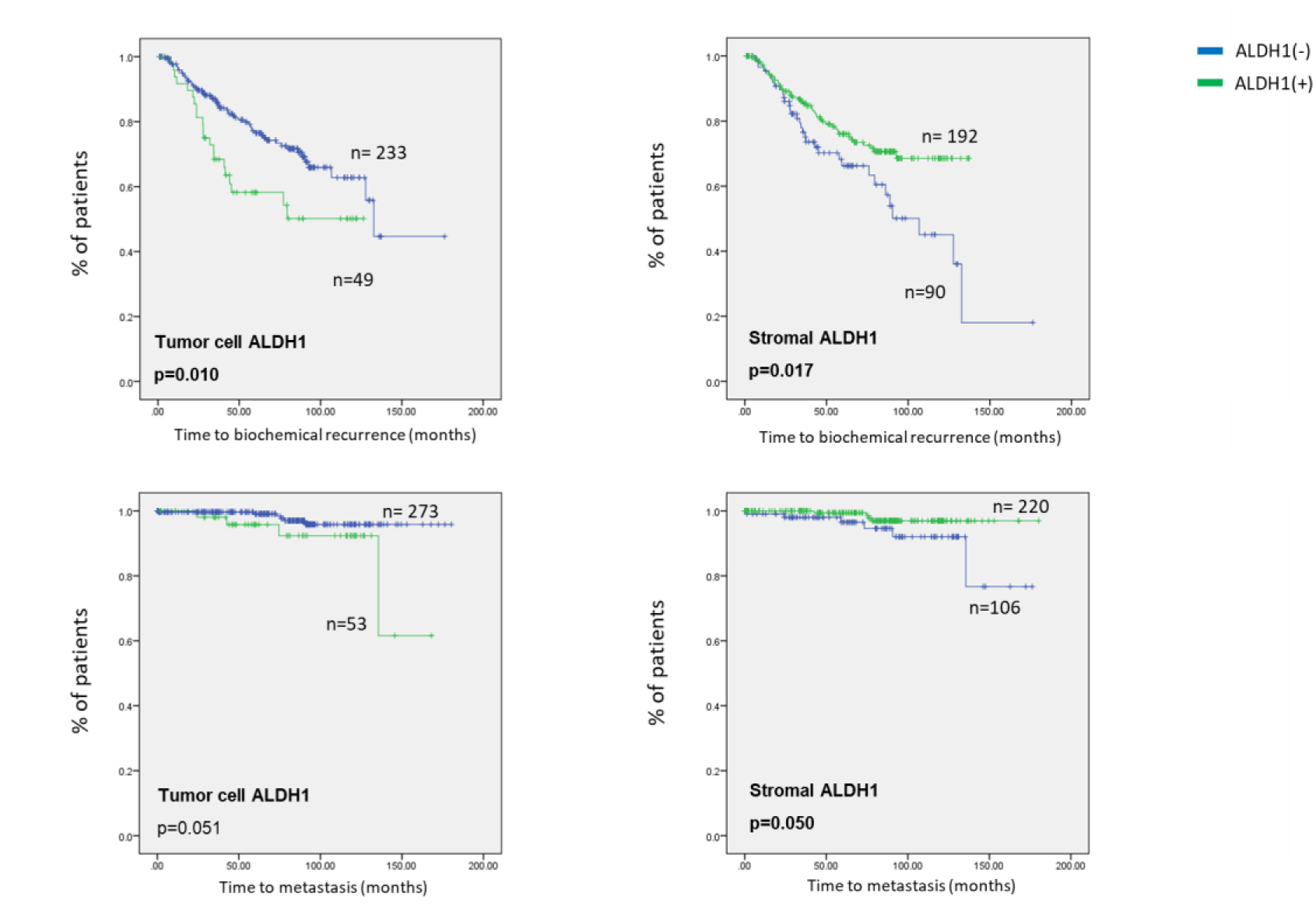
Survival analysis of tumoral and stromal ALDH1 expression in prostate cancer patients. Disease-free survival in prostate cancer patients. Neg indicates negative ALDH1 staining, pos – positive ALDH1 staining

**Table 1.** Clinical relevance of tumoral and stromal ALDH1 expression in prostate cancer patients. Neg indicates negative ALDH1 staining, pos – positive ALDH1 staining, none – no tumoral and stromal ALDH1 staining, S – stromal ALDH1 staining, T+S – tumoral and stromal ALDH1 staining, T – tumoral ALDH1 staining, F – Fisher test otherwise Chi square test.

| Clinical and pathological parameters | Status | ALDH1 stromal |  |  |  | ALDH1 tumoral |  |  |  |
| --- | --- | --- | --- | --- | --- | --- | --- | --- | --- |
|  |  | Neg<br>n | % | Pos<br>n | % | Neg<br>n | % | Pos<br>n | % |
| **Age (years)** | <median | 57 | 50,0 | 127 | 54,3 | 152 | 51,9 | 32 | 58,2 |
|  | >=median | 57 | 50,0 | 107 | 45,7 | 141 | 48,1 | 23 | 41,8 |
|  | total |  |  |  | 0,454 |  |  |  | 0,390 |
| **T status** | T2 | 29 | 25,7 | 92 | 40,2 | 110 | 38,3 | 11 | 20 |
|  | T3-4 | 84 | 74,3 | 137 | 59,8 | 177 | 61,7 | 44 | 80 |
|  | p-value |  |  |  | **0,008** |  |  |  | **0,009** |
| **N status** | N0 | 94 | 85,5 | 214 | 93,4 | 259 | 90,9 | 49 | 90,7 |
|  | N1 | 16 | 14,5 | 15 | 6,6 | 26 | 9,1 | 5 | 9,3 |
|  | p-value |  |  |  | **0,017** |  |  |  | 0,975 |
| **Gleason score sum** | <7 | 46 | 41,1 | 69 | 29,6 | 97 | 33,5 | 18 | 32,7 |
|  | 3+4 | 14 | 12,5 | 61 | 26,2 | 71 | 24,5 | 4 | 7,3 |
|  | 4+3 | 22 | 19,6 | 67 | 28,8 | 72 | 24,8 | 17 | 30,9 |
|  | >7 | 30 | 26,8 | 36 | 15,4 | 50 | 17,2 | 16 | 29,1 |
|  | p-value |  |  |  | **<0.001** |  |  |  | **0,016** |
| **Biochemical recurrence** | no | 57 | 63,3 | 147 | 76,2 | 176 | 75,5 | 28 | 56,0 |
|  | yes | 33 | 36,7 | 46 | 23,8 | 7 | 24,5 | 22 | 44,0 |
|  | p-value |  |  |  | **0,025** |  |  |  | **0,005** |
| **Metastatic relapse** | no | 106 | 94,6 | 229 | 98,3 | 284 | 97,9 | 51 | 92,7 |
|  | yes | 6 | 5,4 | 4 | 1,7 | 6 | 2,1 | 4 | 7,3 |
|  | p-value |  |  |  | 0.084 (F) |  |  |  | 0.058 (F) |
| **Death** | no | 99 | 99,0 | 224 | 99,6 | 273 | 100,0 | 50 | 96,2 |
|  | PCa-related | 1 | 1,0 | 1 | 0,4 | 0 | 0 | 2 | 3,8 |
|  | p-value |  |  |  | 0.521 (F) |  |  |  | **0.025 (F)** |

Positive ALDH1 staining in stromal cells correlated to less aggressive histopathological characteristics of a primary tumor such as lower T (Chi^2^=6.969, p=0.008) and negative N status (Chi^2^=5.717, p=0.017), higher Gleason score (Chi^2^=17.001, p<0.001) as well as less frequently occurring BR (Chi^2^=5.023, p=0.025) (**Tab. 1**). Stromal ALDH1 was also associated with patients’ longer time to BR (Kaplan-Meier log rank analysis, p=0.017) and longer time to metastasis (Kaplan-Meier log rank analysis, p=0.050) (**Fig. 2**). Of note, absence of ALDH1-positive stromal cells (Cox analysis, p=0.050, CI95% 0.846, HR 0.716-1.00) appeared to be also an independent indicator of shorter time to BR in d’Amico high risk PCa patients (n=180) in multivariate analysis (**Tab. 2**).

**Table 2.** Multivariate analysis.

| n=180 |  |  |  |  |  |  |
| --- | --- | --- | --- | --- | --- | --- |
|  | Univariate analysis |  |  | Multivariate analysis |  |  |
|  | p-value | HR | 95% CI for HR | p-value | HR | 95% CI for HR |
| **T status (T3-4 vs. T1-2)** | **0,036** | **4,531** | **1,105-18,579** | 0,073 | 3,673 | 0,885-15,243 |
| **N status (N0 vs. N1)** | 0,261 | 0,559 | 0,203-1,542 | - | - | - |
| **Age (≥64 vs. <64)** | 0,058 | 1,628 | 0,984-2,692 | - | - | - |
| **tumoral ALDH1 pos vs. Neg** | **0,033** | **1,776** | **1,049-3,008** | 0,054 | 1,688 | 0,992-2,872 |
| **stromal ALDH1 neg vs. Pos** | **0,040** | **0,841** | **0,712-0,992** | **0,050** | **0,846** | **0,716-1,000** |

### ALDH1 expression in stromal and tumor cells of prostate cancer lymph node metastases

Since absence of ALDH1 staining in stromal cells of PCa correlated to lymph node positivity, its expression was also examined in lymph node metastases (LNM) of PCa patients (n=98).

Sixty-three of patients were informative for ALDH1 staining in LNM samples. Twenty-one (33.3%) and 26 (41.3%) of those patients were positive for ALDH1 in tumor and stromal cells, respectively. The number of patients with ALDH1-positive tumor cells was significantly higher when compared to primary tumors (33.3% vs. 15.8%, Ch^2^=10.874, p=0.001). On the contrary, the number of cases with ALDH1-positive stroma was significantly lower in LNMs than in primary tumors (41.3% vs. 68.1%, Ch^2^=16.710, p<0.001). The mean percentage of ALDH1-positive tumor cells per LNM sample was 11% and tended to be higher than the mean percentage in a primary tumor (two-samples t-test, p=0.052).

The comparison of 13 matched pairs of primary tumor and LNMs revealed that 8 (61.5%) cases had the same status of ALDH1 in tumor cells both in primary tumor and LNM, whereas 4 (30.8%) patients had ALDH1-positive tumor cells only in LNM (**Suppl. Tab. 2**). Eight (61.6%) pairs of primary tumor and LNM displayed identical ALDH1 staining in stromal cells in both sites, whereas 2 (15.4%) patients had ALDH1-positive stromal cells exclusively in LNM (**Suppl. Tab. 2**). Tumor cell and stromal ALDH1 staining in LNM did not correlate to any clinico-pathological parameter or survival (data not shown).

### ALDH1 expression in stromal and tumor cells of castration resistant prostate cancer metastases

To complete the picture of ALDH1 in PCa progression, we investigated its presence in tumor and stromal cells of CRPC metastases to distant organs in an independent cohort of unmatched metastatic castrate-resistant patients (n=21). Twenty-one and 18 patients were informative for ALDH1 staining in bone metastases or metastases to other distant organs (liver, lung, kidney), respectively.

ALDH1-positive tumor cells were found in at least one distant organ in 17 (81.0%) patients: 11 (52.4%) and 16 (88.9%) of bone and visceral metastases, respectively. ALDH1-positive tumor cells seemed to be more frequent in metastases to other organs than in metastases to bones (Fisher’s exact test, p=0.0327). The mean percentage of ALDH1-positive tumor cells was 20.6% (n=40) in metastases to bones and 30.8% (n=36) in metastases to other distant organs. ALDH1 staining in tumor cells appeared significantly more frequently in metastases than in primary tumors (Chi-square test, both p<0.001) and the percentage of ALDH1-positive tumor cells per sample was higher when compared to primary tumors (independent samples t-test, p=0.0125 and p<0.001, respectively).

None of the patients had ALDH1(+) stroma in both metastatic sites. There was also no difference in frequency of ALDH1-positive stromal cells between metastases to bones or other distant organs: they were found in 5 (23.8%) and 5 (27.8%) of bone and visceral metastases, respectively. ALDH1 staining in stromal cells appeared significantly less frequently in metastases than in primary tumors (Chi-square test, both p<0.001).

Within bone metastases, 38.1% of patients were negative for ALDH1 both in tumor cells and stroma. ALDH1 staining in exclusively tumor cells was found in 38.1% of patients, only in stroma – 9.5%, whereas in both tumor and stromal cells - in 14.3% of cases. In metastases to other organs (liver, lung and kidney), 27.8% patients were positive for ALDH1 in tumor and stromal cells, and 11.15% were negative for ALDH1 in both compartments. Interestingly, none of the patients were positive for ALDH1 staining detected exclusively in stromal cells of these metastases, whereas 72.2% were positive for ALDH1 detected only in tumor cells.

### Molecular characterization of primary prostate cancers in context of ALDH1 expression in tumor and stromal cells

To better characterize the putative correlation of ALDH1 staining in tumor cells and the stromal compartment of PCa with tumor cell differentiation and aggressiveness, ALDH1 expression was compared to selected molecular markers associated with proliferation (Ki-67), de-differentiation of tumor cells (Gleason score, luminal cytokeratins (CK) CK8/18 and CK19, basal cytokeratins CK5/6 and CK14) as well as EMT-related markers (E-cadherin, N-cadherin, vimentin) and epithelial cell marker EpCAM (**Tab. 3**; ref: 18). Of note, in order to better follow the correlations between molecular factors and evaluated ALDH1 staining, this comparative analysis was performed between individual tumor samples, not between patients.

**Table 3.** Molecular characteristics of prostate cancers in comparison to ALDH1 expression in tumor and stromal cells. neg indicates negative ALDH1 staining, pos – positive ALDH1 staining, CK – cytokeratin

| Molecular marker |  | Total cohort |  | ALDH1 stromal |  |  |  | ALDH1 tumoral |  |  |  |
| --- | --- | --- | --- | --- | --- | --- | --- | --- | --- | --- | --- |
|  |  | n | % | Neg |  | Pos |  | Neg |  | Pos |  |
|  |  |  |  | n | % | n | % | n | % | n | % |
| Ki-67 | neg | 608 | 86,4 | 114 | 85,1 | 337 | 83,6 | 391 | 83,2 | 60 | 89,6 |
|  | pos | 96 | 13,6 | 20 | 14,9 | 66 | 16,4 | 79 | 16,8 | 7 | 10,4 |
|  |  | 704 |  | p=0,691 |  |  |  | p=0,184 |  |  |  |
| CK8/18 | neg | 281 | 41,1 | 61 | 47,3 | 145 | 36,7 | 185 | 40,5 | 21 | 31,3 |
|  | pos | 402 | 58,9 | 68 | 52,7 | 250 | 63,3 | 272 | 59,5 | 46 | 68,7 |
|  |  | 683 |  | p=0,033 |  |  |  | p=0,153 |  |  |  |
| CK19 | neg | 361 | 56,7 | 75 | 62,0 | 203 | 54,4 | 242 | 56,3 | 36 | 56,3 |
|  | pos | 276 | 43,3 | 46 | 38,0 | 170 | 45,6 | 188 | 43,7 | 28 | 43,8 |
|  |  | 637 |  | p=0,145 |  |  |  | p=0,997 |  |  |  |
| CK5/6 | neg | 625 | 94,8 | 122 | 93,8 | 361 | 95,8 | 420 | 9,2 | 63 | 95,5 |
|  | pos | 34 | 5,2 | 8 | 6,2 | 16 | 4,2 | 21 | 4,8 | 3 | 4,5 |
|  |  | 659 |  | p=0,377 |  |  |  | p=0,938 |  |  |  |
| CK14 | neg | 504 | 78,4 | 96 | 79,3 | 291 | 77,6 | 347 | 79,6 | 40 | 66,7 |
|  | pos | 139 | 21,6 | 25 | 20,7 | 84 | 22,4 | 89 | 20,4 | 30 | 33,3 |
|  |  | 643 |  | p=0,688 |  |  |  | p=0,023 |  |  |  |
| E-cad | neg | 279 | 43,6 | 3 | 45,7 | 154 | 40,6 | 183 | 42,4 | 24 | 38,1 |
|  | pos | 361 | 56,4 | 63 | 54,3 | 225 | 59,4 | 249 | 57,6 | 39 | 61,9 |
|  |  | 640 |  | p=0,334 |  |  |  | p=0,521 |  |  |  |
| N-cad | neg | 429 | 66,5 | 89 | 70,1 | 253 | 65,7 | 300 | 67,3 | 42 | 63,6 |
|  | pos | 216 | 33,5 | 38 | 29,9 | 132 | 34,3 | 146 | 32,7 | 24 | 36,4 |
|  |  | 645 |  | p=0,365 |  |  |  | p=0,559 |  |  |  |
| Vimentin | neg | 367 | 54,1 | 78 | 61,4 | 286 | 73,3 | 320 | 70,8 | 44 | 7,7 |
|  | pos | 311 | 45,9 | 49 | 38,6 | 104 | 26,7 | 132 | 29,2 | 21 | 32,3 |
|  |  | 678 |  | p=0,011 |  |  |  | p=0,608 |  |  |  |
| EpCAM | neg | 205 | 37,1 | 51 | 60,0 | 99 | 30,4 | 134 | 36,0 | 16 | 41,0 |
|  | pos | 347 | 62,9 | 34 | 40,0 | 227 | 69,6 | 238 | 64,0 | 23 | 59,0 |
|  |  | 552 |  | p<0.001 |  |  |  | p=0,537 |  |  |  |
| EMT | CK8/18(+)Vim(-) | 245 | 38,4 | 34 | 28,3 | 160 | 42,9 | 163 | 38,0 | 31 | 48,4 |
|  | CK8/18(-)Vim(-) | 196 | 30,7 | 38 | 31,7 | 113 | 30,3 | 139 | 32,4 | 12 | 18,8 |
|  | CK8/18(+)Vim(+) | 135 | 21,2 | 29 | 24,2 | 75 | 20,1 | 91 | 21,2 | 13 | 20,3 |
|  | CK8/18(-)Vim(+) | 62 | 9,7 | 19 | 15,8 | 25 | 6,7 | 36 | 8,4 | 8 | 12,5 |
|  |  | 638 |  | p=0,003 |  |  |  | p=0,111 |  |  |  |

ALDH1 expression in tumor cells was significantly associated with expression of basal cytokeratin CK14 (Chi^2^=4.786, p=0.029) and a higher Gleason score determined in individual tumors (Chi^2^=14.666, p=0.002) (**Tab. 3**).

On the contrary, positive stromal expression of ALDH1 was associated with a lower Gleason score determined in individual tumors (Chi^2^=14.429, p=0.002) (**Tab. 3**). Stromal ALDH1 expression was also present in tumors, which expressed significantly more frequently luminal CK8/18 (Chi^2^=4.561, p=0.033) and epithelial cell marker EpCAM (Chi^2^=25.543, p<0.001), and displayed rarely EMT-like characteristics defined by CK8/18(-)Vimentin(+) (Chi^2^=14.139, p=0.003) (**Tab. 3**).

## Discussion

It the current study, for the first time, ALDH1 was evaluated as a protein of potential dual clinical relevance in PCa. ALDH1 staining in tumor cells was associated with tumor aggressiveness and disease progression, whereas ALDH1 staining in stromal cells was shown to predict better patients’ outcome and corresponded to less aggressive clinical and molecular characteristics of the analyzed tumors. ALDH1-positive tumor and stromal cells were also, for the first time, demonstrated to be present in hormone-naïve PCa metastases to lymph nodes and CRPC metastases to distant organs.

Tumor cell ALDH1 expression was assessed in 16% of patients. The percentages of ALDH1-positve prostate tumor cells showed a broad range and a low mean as demonstrated also in breast cancer (3; 20-23). The correlations of tumor cell ALDH1 expression to a more aggressive course of disease such as higher T status, and biochemical recurrence, as well as shorter time to BR and metastasis support the hypothesis that ALDH1-positive tumor cells might be indeed the subpopulation of cells involved in tumor progression. Additionally, the presence of ALDH1 in tumor cells was associated with a less differentiated tumor epithelial phenotype (higher Gleason score) and expression of basal cytokeratin CK14. PCa stem cells are believed to originate from poorly differentiated basal cells (24). Therefore, these associations might corroborate with putative stem-cell-like status of the evaluated ALDH1-positive tumor cells. Taken together, the collected data might explain correlations between presence of ALDH1-positive tumor cells and tumor progression.

The stromal ALDH1 staining has not yet been examined in PCa. In the present study, stromal ALDH1 staining was detected in 67% of primary PCa and appeared to be associated with both a less aggressive tumor cell phenotype and more favorable outcome. ALDH1-positive stroma correlated inversely to T and N status, Gleason score sum, as well as BR. Stromal positivity also indicated a longer time to BR and metastasis, and appeared even to be an independent predictor of good prognosis in the subgroup of d’Amico high risk patients. It is reasonable to conclude that ALDH1-positive stromal cells might play a protective role in PCa, which is similar to the observation in breast cancer (2).

ALDH1-positive stromal cells might hypothetically secrete retinoic acid and consequently in this way suppress tumor aggressiveness (2). Retinoic acid is known to induce tumor cell differentiation and inhibit tumor cell proliferation and migration (7-9). Accordingly, the presence of ALDH1-positive stromal cells in our cohort of PCa’s was associated with more differentiated tumor cell phenotype (presence of luminal CK8/18, lower Gleason score). Additionally, ALDH1 stromal staining correlated with a higher expression of EpCAM and less frequent signatures of EMT in tumor cells indicating a more epithelial phenotype and rather limited migration abilities.

To the best of our knowledge, the current study is the first one evaluating ALDH1 expression in both tumor and stromal cells in PCa metastases to lymph nodes and distant organs.

Of note, the number of patients with ALDH1-positive tumor cells in LNMs or distant metastases was significantly higher than those positive for tumor cell ALDH1 in primary tumors. Additionally, the mean percentage of ALDH1-positive tumor cells per LNM or distant metastasis sample was higher than in primary tumors. The higher frequency of ALDH1-positive tumor cells among LNM and distant metastases might suggest that ALDH1-positive PCa cells have an increased potential to disseminate to distant organs and/or produce overt metastasis. It might also indicate that an aggressive phenotype of PCa is easier to be identified in LNM than in primary tumor. This hypothesis might have important impact on diagnostics, meaning that LNMs biopsies should be taken and analysed in the context of specific markers in order to better define PCa patients at the high risk of progression. It also supports the idea that further biopsies of tumor at secondary sites should be done in order to tailor the treatment once the overt metastasis occur.

Tumor and stromal cells interact with each other, which might influence tumor development at different sites. Interestingly, significantly fewer patients were positive for ALDH1 stromal staining in LNMs and distant metastasis than in primary tumors. It might be concluded that ALDH1-positive immune cells that suppress tumor outgrowth, could infiltrate tumors less frequently at secondary sites than at primary sites. Alternatively, it might be assumed that at least some tumor cells develop the efficient mechanisms to avoid immune response and only those are able to establish overt metastasis. This observation merits further investigation.

## Conclusions

The presented data support the hypothesis that ALDH1 might play an opposing role in tumor cells and tumor-associated stromal cells in PCa (**Fig. 3**). We hypothesize that ALDH1-positive stromal cells can induce differentiation and attenuate PCa progression, whereas ALDH1 expressed in tumor cells indicates a more aggressive phenotype and worse clinical outcome. This observation is similar to previous reports in breast cancer patients (1, 2). Therefore, we hypothesize the presence of ALDH1(+) stromal cells might also attenuate the growth and progression of other epithelial malignancies.

**Figure 3.**
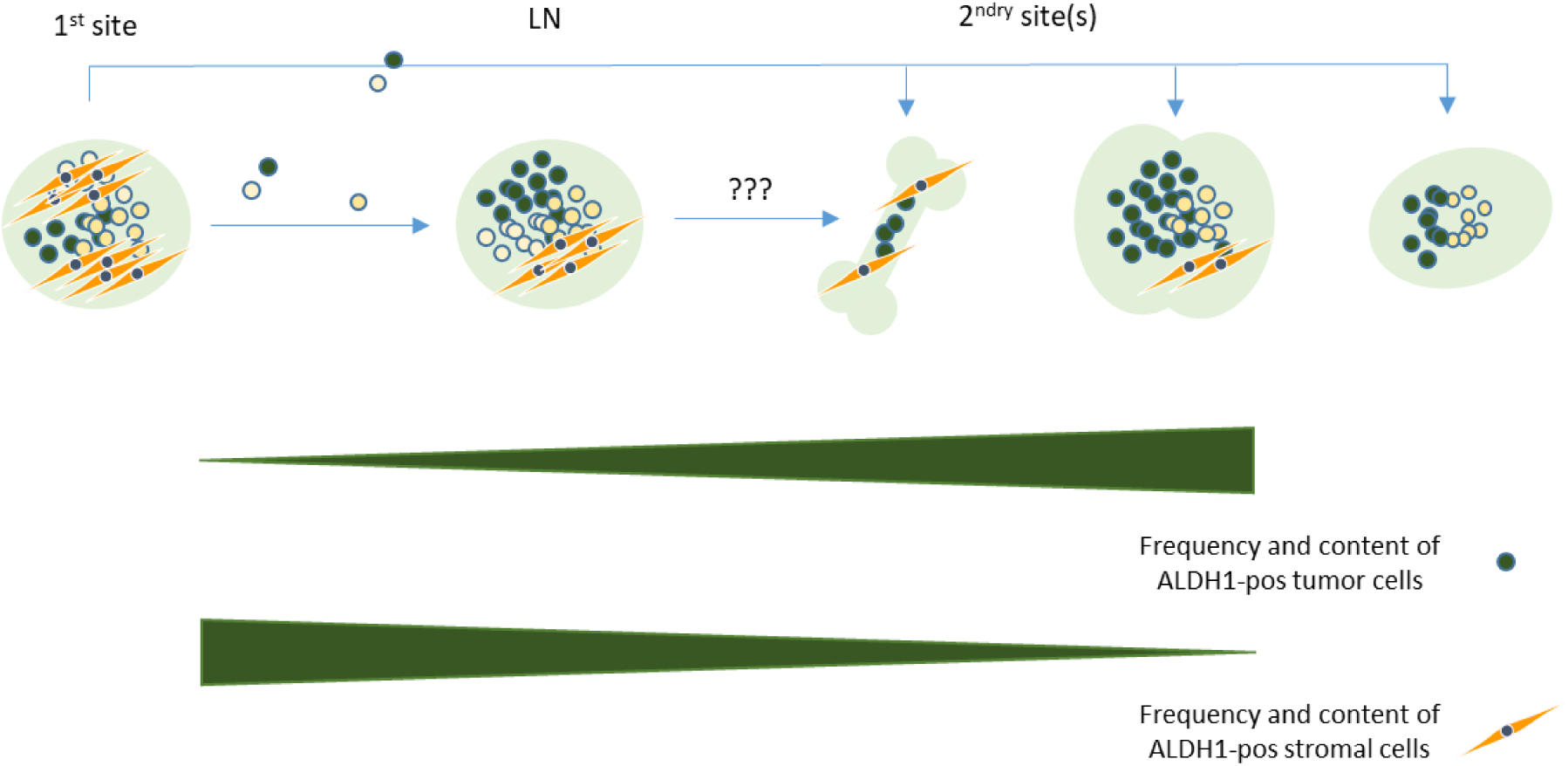
Model of hypothetical involvement of ALDH1 in tumor and stromal cells in tumor progression.

## Acknowledgements

We thank the patients and their families who were willing to participate in the Prostate Cancer Donor Program, the investigators Drs. Robert Vessella, Celestia Higano, Bruce Montgomery, Evan Yu, Peter Nelson, Paul Lange, Martine Roudier, and Lawrence True for their contributions to the University of Washington Medical Center Prostate Cancer Donor Rapid Autopsy Program as well as Małgorzata Stoupiec for excellent technical assistance.

This research was supported by funding by the Movember GAP1 Funding Award: Integrated Global CTCs Biomarker Project (Movember Foundation, Australia) (K.P.), the ERC Advanced Investigator Grant DISSECT (K.P.), the Pacific Northwest Prostate Cancer SPORE (P50CA97186), the PO1 NIH grant (PO1CA085859) and the Richard M. LUCAS Foundation.

## List of Supplementary Tables

**Supplementary Table 1.** Clinico-pathological data for prostate cancer patients included in the study. Note that due to the missing values not all numbers sum up to 398 cases DFS indicates disease free survival

| Clinical and pathological parameters | Status | n | % |
| --- | --- | --- | --- |
| **Age (years)** | median | 64 |  |
|  | range | 46-77 |  |
|  | <median | 207 | 52,0 |
|  | >=median | 191 | 48,0 |
|  | total | 398 |  |
| **T status** | T2 | 152 | 38,2 |
|  | T3 | 225 | 56,5 |
|  | T4 | 21 | 5,3 |
|  | total | 398 |  |
| **N status** | N0 | 355 | 93,7 |
|  | N1 | 20 | 5,3 |
|  | N2 | 4 | 1,1 |
|  | total | 379 |  |
| **M status** | M0 | 391 | 100 |
|  | M1 | 0 | 0 |
|  | total | 391 |  |
| **Gleason score** | <7 | 135 | 34,1 |
|  | 3+4 | 93 | 23,5 |
|  | 4+3 | 94 | 23,7 |
|  | >7 | 74 | 18,7 |
|  | total | 396 |  |
| **Biochemical recurrence** | no | 235 | 72,5 |
|  | yes | 89 | 27,5 |
|  | total | 324 |  |
| **Metastatic relapse** | no | 383 | 97,0 |
|  | yes | 12 | 3,0 |
|  | total | 395 |  |
| **Death** | no | 370 | 93,4 |
|  | PCa-related | 2 | 0,5 |
|  | PCa-unrelated | 11 | 2,8 |
|  | unknown | 13 | 3,3 |
|  | total | 396 |  |

**Supplementary Table 2.** ALDH1 expression in matched pairs of prostate cancer primary tumors and lymph node metastases.

| PT | LNM | Tumor cells |  | Stromal cells |  |
| --- | --- | --- | --- | --- | --- |
|  |  | n | % | n | % |
| neg | neg | 7 | 53,8 | 4 | 30,8 |
| neg | pos | 4 | 30,8 | 2 | 15,4 |
| pos | pos | 1 | 7,7 | 4 | 30,8 |
| pos | neg | 1 | 7,7 | 3 | 23,1 |
|  | total | 13 |  | 13 |  |

## References

1. Resetkova E, Reis-Filho JS, Jain RK, et al. Prognostic impact of ALDH1 in breast cancer: a story of stem cells and tumor microenvironment. Breast Cancer Res Treat 2010;123:97–108.

2. Bednarz-Knoll N, Nastały P, Żaczek A, et al. -Stromal expression of ALDH1 in human breast carcinomas indicates reduced tumor progression—Oncotarget, 2015;6:26789–803.

3. Ginestier C, Hur ME, Charafe-Jauffret E, et al. ALDH1 Is a Marker of Normal and Malignant Human Mammary Stem Cells and a Predictor of Poor Clinical Outcome. Cell Stem Cell 2007;1:555–67.

4. Ma I, Allan AL. The role of human aldehyde dehydrogenase in normal and cancer stem cells. Stem Cell Rev. 2011;7: 292–306.

5. Tirino V, Desiderio V, Paino F, et al. Cancer stem cells in solid tumors: an overview and new approaches for their isolation and characterization. FASEB J. 2013;27:13–24.

6. Kam RK, Deng Y, Chen Y, Zhao H. Retinoic acid synthesis and functions in early embryonic development. Cell Biosci. 2012;2:11.

7. Ginestier C, Wicinski J, Cervera N, et al. Retinoid signaling regulates breast cancer stem cell differentiation. Cell Cycle 2009;8:3297–302.

8. Tang XH, Gudas LJ. Retinoids, retinoic acid receptors, and cancer. Annu Rev Pathol 2011;6: 345–64.

9. Guan J, Zhang H, Wen Z, et al. Retinoic acid inhibits pancreatic cancer cell migration and EMT through the downregulation of IL-6 in cancer associated fibroblast cells. Cancer Lett 2014;345:132–9.

10. Honeth G, Lombardi S, Ginestier C, et al. Aldehyde dehydrogenase and estrogen receptor define a hierarchy of cellular differentiation in the normal human mammary epithelium. Breast Cancer Res 2014;16:R52.

11. Liu Y, Lv DL, Duan JJ, et al. ALDH1A1 expression correlates with clinicopathologic features and poor prognosis of breast cancer patients: a systematic review and meta-analysis. BMC Cancer. 2014, 17; 14:444.

12. Doherty RE, Haywood-Small SL, Sisley K, Cross NA. Aldehyde dehydrogenase activity selects for the holoclone phenotype in prostate cancer cells. Biochem Biophys Res Commun 2011;414:801–7.

13. Li T, Su Y, Mei Y, et al. ALDH1A1 is a marker for malignant prostate stem cells and predictor of prostate cancer patients’ outcome. Lab Invest 2010;90:234–44.

14. Le Magnen C, Bubendorf L, Rentsch CA, et al. Characterization and clinical relevance of ALDHbright populations in prostate cancer. Clin Cancer Res 2013;19:5361–71.

15. van den Hoogen C, van der Horst G, Cheung H, et al. High aldehyde dehydrogenase activity identifies tumor-initiating and metastasis-initiating cells in human prostate cancer. Cancer Res 2010;70:5163–73.

16. Cojoc M, Peitzsch C, Kurth I, et al. Aldehyde Dehydrogenase Is Regulated by β-Catenin/TCF and Promotes Radioresistance in Prostate Cancer Progenitor Cells. Cancer Res 2015;75:1482–94.

17. Bednarz N, Eltze E, Semjonow A, et al. BRCA1 loss in small subpopulations of tumor cells indicates an increased risk of prostate cancer spread and early biochemical recurrence. Clin Cancer Res 2010;16:3340–8.

18. Omari A, Nastały P, Bałabas A, et al.-Somatic aberrations of BRCA1 gene are associated with progressive and stem cell-like phenotype of prostate cancer – bioRxiv, 2018, doi.org/10.1101/271312

19. McShane LM, Altman DG, Sauerbrei W, et al. REporting recommendations for tumor MARKer prognostic studies (REMARK). Breast Cancer Res Treat 2006;100:229–35.

20. Charafe-Jauffret E, Ginestier C, Iovino F, et al. Aldehyde dehydrogenase 1-positive cancer stem cells mediate metastasis and poor clinical outcome in inflammatory breast cancer. Clin Cancer Res 2010;1:45–55.

21. Deng S, Yang X, Lassus H, et al. Distinct expression levels and patterns of stem cell marker, aldehyde dehydrogenase isoform 1 (ALDH1), in human epithelial cancers. PLoS One 2010;21:5.

22. Gong C, Yao H, Liu Q, et al. Markers of tumor-initiating cells predict chemoresistance in breast cancer. PLoS One 2010;20:15630.

23. Yoshioka T, Umekita Y, Ohi Y, et al. Aldehyde dehydrogenase 1 expression is a predictor of poor prognosis in node-positive breast cancers: a long-term follow-up study. Histopathology. 2011;58:608–16.

24. Smith BA, Sokolov A, Uzunangelov V et al. A basal stem cell signature identifies aggressive prostate cancer phenotypes. Proc Natl Acad Sci U S A. 2015 112(47):E6544–52

